# Long-term evolution experiments fully reveal the potential for thermal adaptation

**DOI:** 10.1101/2024.02.15.580572

**Authors:** Marta A. Antunes, Afonso Grandela, Margarida Matos, Pedro Simões

## Abstract

Global warming is leading to worldwide biodiversity decline at a fast pace. Evolutionary responses may be crucial in allowing organisms to cope with prolonged effects of climate change. This urges the need for a better understanding of the dynamics of adaptation to warming environments. In particular, addressing how reproductive success evolves in deteriorating environments is extremely relevant, as this trait is more likely constrained at lower temperatures than upper physiological thermal limits. Experimental evolution under a warming environment can elucidate the potential of populations to respond to rapid environmental changes. The few studies following such framework lack analysis of long-term response. We here focus on the long-term thermal evolution of two *Drosophila subobscura* populations, from different European latitudes, under warming temperatures. We estimated the reproductive success of these populations in two test environments: the ancestral (control) and the warming environment after 39 and 52 generations of thermal evolution. We found that a relevant long-term adaptive response to warming temperatures can occur, but the pace of such response is slow. In addition, we observed contrasting responses in the ancestral environmental and differences in the evolutionary dynamics between populations of distinct histories, with those originally from higher latitude only showing an adaptive response under the to the warming environment regime in at a later generation. This study reinforces the need for long-term evolution experiments to fully reveal the potential for thermal response. It also highlights that the scrutiny of several populations in this context is needed for a measure of variation within a species. Accounting for these sources of variation - both temporal and spatial - will allow for more robust assessments of climate change responses.

## Introduction

Climate change is causing declines in biodiversity at an unprecedented pace and is likely to continue doing so in the future (IPCC 2023). Evolution may play an important role in enhancing organisms’ ability to cope with the prolonged effects of climate change (Urban et al. 2016). It is thus crucial to understand the dynamics of adaptation to sustained warming temperatures to better predict the impacts of climate change (Martin et al. 2023; Edelsparre et al. 2024). In particular, addressing how reproductive success evolves in warmer environments is extremely relevant, as it has been shown that negative effects of heat stress on fertility occur quite often at lower temperatures than physiological impairment (Walsh et al. 2019a,b; Parratt et al. 2021; van Heerwaarden and Sgrò 2021). In fact, fertility patterns have been shown to track species’ geographical distribution in nature, highlighting its role as a predictor of biodiversity changes due to climate change (Parratt et al. 2021; van Heerwaarden and Sgrò 2021).

The possibility of an adaptive response to a deteriorating environment is shaped by two main factors: the standing genetic variation within populations and the rate of environmental change (Burger and Lynch 1995; Bell 2017). The amount of genetic variation sets the limit to the rate of environmental change a given population can sustain. If genetic variation is present and provided that the environmental change is not too high, the average phenotype may evolve in parallel though lagging behind the changing optimum. In contrast, when genetic variation for fitness-related traits is low and/or evolutionary constraints between traits occur, the rates of evolution will be limited and populations may not be able to keep pace with environmental change (Burger and Lynch 1995). Low population size will expectedly compromise genetic variation and as such will hinder the effectiveness of evolution (Gomulkiewicz and Holt 1995; Martin et al. 2023). Under such demographic scenario, adaptation to the new conditions may only be possible if the rate of environment change is very slow or ceases (Hoffmann and Sgró 2011; Lindsey et al. 2013; Bell 2017).

Experimental evolution is a powerful tool to deepen our understanding about the tempo and mode of thermal evolution, as it allows to follow the real-time evolution of populations under relevant thermal settings (Kawecki et al. 2012; Rogell et al. 2014; Kellermann and van Heerwaarden 2019; Santos et al. 2021; van Heerwaarden and Sgrò 2021). Furthermore, it allows to directly test the role of specific environmental factors, e.g. such as temperature, while confounding environmental effects are minimized relative to field studies. Some studies in ectotherms (mostly Drosophila) have focused on evolutionary responses of reproductive traits to experimental warming but most have covered a limited temporal range – around 20 generations or less (Schou et al. 2014; Kinzner et al. 2019; van Heerwaarden and Sgrò 2021). In general, these studies have found limited evolutionary response to warming in reproductive traits (Schou et al. 2014; Kinzner et al. 2019; van Heerwaarden and Sgrò 2021). On the other hand, Rogell et al. (2014) found an evolution of increased reproductive performance in response to incremental warming (rate of 0.3 C per generation) during 18 generations, with populations of seed beetle *Callosobruchus maculatus* evolving under higher temperatures presenting increased offspring production relative to their controls. Importantly, this effect on fecundity was only observed 23 generations after the incremental setting (Rogell et al. 2014) and not immediately after the increment (Hallsson and Björklund 2012). These experiments suggest that the tempo of adaptive change to new thermal conditions is likely slow – possibly due to limited genetic variation for reproductive traits - and that a rapid rate of environmental change poses serious problems to a timely adaptive response. In addition, reductions in genetic variation might result from prolonged evolution under stressful conditions due to the continuous effects of genetic drift when effective population size is rather low and/or there are population crashes causing bottlenecks. Considering these different sources of genetic variation, long-term responses to incremental temperatures may be highly variable between populations and species. One caveat of the thermal experimental evolution studies reported above is that they neglect effects of variation at the inter-population level - namely by studying different geographical populations of a species - and how it affects the adaptive response.

Experimental evolution studies under controlled conditions offer the possibility to assess adaptation to specific conditions as well as to detect associated costs of adaptation by testing control and experimental populations in both the ancestral and a novel environment during real-time evolution, something that is not straightforward in field studies (though this may be achieved through reciprocal transplant experiments, Kawecki and Ebert 2004). Adaptation to a specific thermal environment can lead to a reduced performance in the ancestral (thermal) environment, for instance, due to the expression of genes with antagonistic pleiotropy (Kawecki and Ebert 2004). However, detection of costs of adaptation in general has been evasive possibly because of their low magnitude (Hereford 2009; Bono et al. 2017).

*Drosophila subobscura* is a species that shows evolutionary responses to climatic factors with repeatable clines of body size (Huey et al. 2000; Gilchrist et al. 2004) and chromosomal inversions (Prevosti et al. 1988). The inversion polymorphism in this species is particularly relevant as it tracks global warming (Balanyà et al. 2006; Rezende et al. 2010). This species also presents high levels of thermal plasticity for reproductive traits(Fragata et al. 2016; Simões et al. 2020; Santos et al. 2021). Our team has been studying how populations of this species evolve under an increasingly warmer environment (Santos et al. 2021). We have been focusing on populations originally from northern (high latitude) and southern (low latitude) European locations and maintained as separate populations for 70 generations in a benign laboratory environment, to address the relevance of different historical backgrounds during thermal evolution (Santos et al. 2021, 2023a,b). Prior to the imposition of the thermal selection regime, in agreement with expectations, these populations showed evidence of fast adaptation to the lab environment during around 30 generations of laboratory evolution - the stronger phase of lab adaptation (Simões et al. 2017). Despite this clear phenotypic response, karyotypic and genomic differentiation resulting from the distinct historical origins was maintained in that period (Simões et al. 2017; Barreira 2023).

To study thermal adaptation in these populations, we applied an incremental warming regime (0.2°C increase per generation) until generation 24 and then the thermal environment was kept the same across generations until the end of the experiment, in a daily fluctuating regime spanning both high and low temperatures (with daily peaks of respectively 11 C and 5 C above and below 18 C, the control conditions). Given the fact that this warming regime includes daily fluctuations and considering the steeper decline in fitness towards upper thermal limits - typical of thermal performance curves – this will likely lead to a higher impact of high temperature on organism performance (i.e. due to the expected effect of Jensen’s inequality, see (Martin and Huey 2008; Colinet et al. 2015). This effect would enhance selective pressures towards adaptation to the high temperatures, contributing to a selective response to warming. We previously found that the evolutionary response of populations to warming conditions was slow and specific to the history of the populations. In fact, thermal adaptation was only observed in the populations of low latitude, and only after 39 generations, while neither population showed signs of an evolutionary response after 22 generations in the warming environment (Santos et al. 2023b). We also found minor evidence supporting costs for thermal adaptation in the populations of southern origin (Santos et al. 2023b). It remains to be tested whether longer-term evolution would allow for an adaptive response in the higher latitude populations. These findings call for a better understanding of the evolutionary dynamics of thermal responses that can occur in different populations.

We here focus on the long-term adaptive response of our warming-evolved populations by analysing their reproductive success after more than 50 generations of thermal evolution. Additionally, we aim to test the dynamics of thermal adaptation, comparing the response to that of generation 39 (Santos et al. 2023b). Finally, coupling data from both long-term assays (generations 39 and 52) we allow for a more robust analysis of the adaptive response. Specifically, with such long term analysis we expect that: 1) the fact that the warming regime was kept unchanged for more than 25 generations by the time the last assay was conducted will have facilitated the adaptive response in both high and low latitude populations; 2) by assaying populations in the ancestral and the warming environment we expect a stronger response in the warming environment and eventually adaptive costs expressed by a poor performance of warming populations in the control environment.

## Material and methods

### Population maintenance and thermal selection regimes

The experimental populations used in this study were obtained from two collections of natural populations of *Drosophila subobscura*, done in August/September 2013 in Adraga, Portugal (38°48°N) and Groningen, The Netherlands (53° 13°N). 213 founder females were collected in Adraga and 170 in Groningen (Simões et al. 2017). These collections gave rise to the PT and NL laboratorial populations. In the two first generations in the lab, females were maintained as families to avoid losing genetic variability due to sampling effects in the initial generations of lab foundation (see Fragata et al. 2014, cf. Santos et al. 2013). Outbred populations were originated in generation three as described in Fragata et al. 2014. These populations were maintained in 30 cm3 glass vials with controlled densities in eggs (70 eggs per vial) and adults (40 adults per vial), with discrete generations of 28 days, 12L:12D photoperiod and a constant 18°C temperature. 18°C is a benign temperature for this species (e.g. see Santos et al. 2005, Fragata et al. 2014) so we do not expect that temperature *per se* is a major factor in determining patterns of lab adaptation. At the start of each generation, eggs laid by adults during a 24h period were counted and allocated to developmental vials (24 vials per replicate population). Upon emergence, imagoes were collected from each development vial during four days. These individuals were subsequently mixed together with CO_2_ anaesthesia, redistributed as adults in 24 vials per replicate population and maintained for one week. After that, egg collection for the new generation took place. As such, adults were seven to ten days old. By generation four in the lab, three replicate populations were created of each founding population, leading to PT1-3 and NL1-3 populations.

This maintenance protocol was maintained for 70 generations, after which a new thermal selection regime was created. The derivation of the new thermal regime was done by dividing the egg collection of each ancestral PT and NL replicate population in equal parts, that were assigned to the two different temperature regimes: the control with the standard maintenance protocol, at constant 18°C) and the warming regime (the new thermal regime, see Figure 1 and below). For example, the NL1 replicate population generated the control NL1 and the warming WNL1 populations. PT and NL population, used as control populations in the experiment, are assumed to represent the ancestral state of the WPT and WNL populations, being maintained as much as possible in synchrony with the warming populations (see also Figure 1).

**Figure 1.**
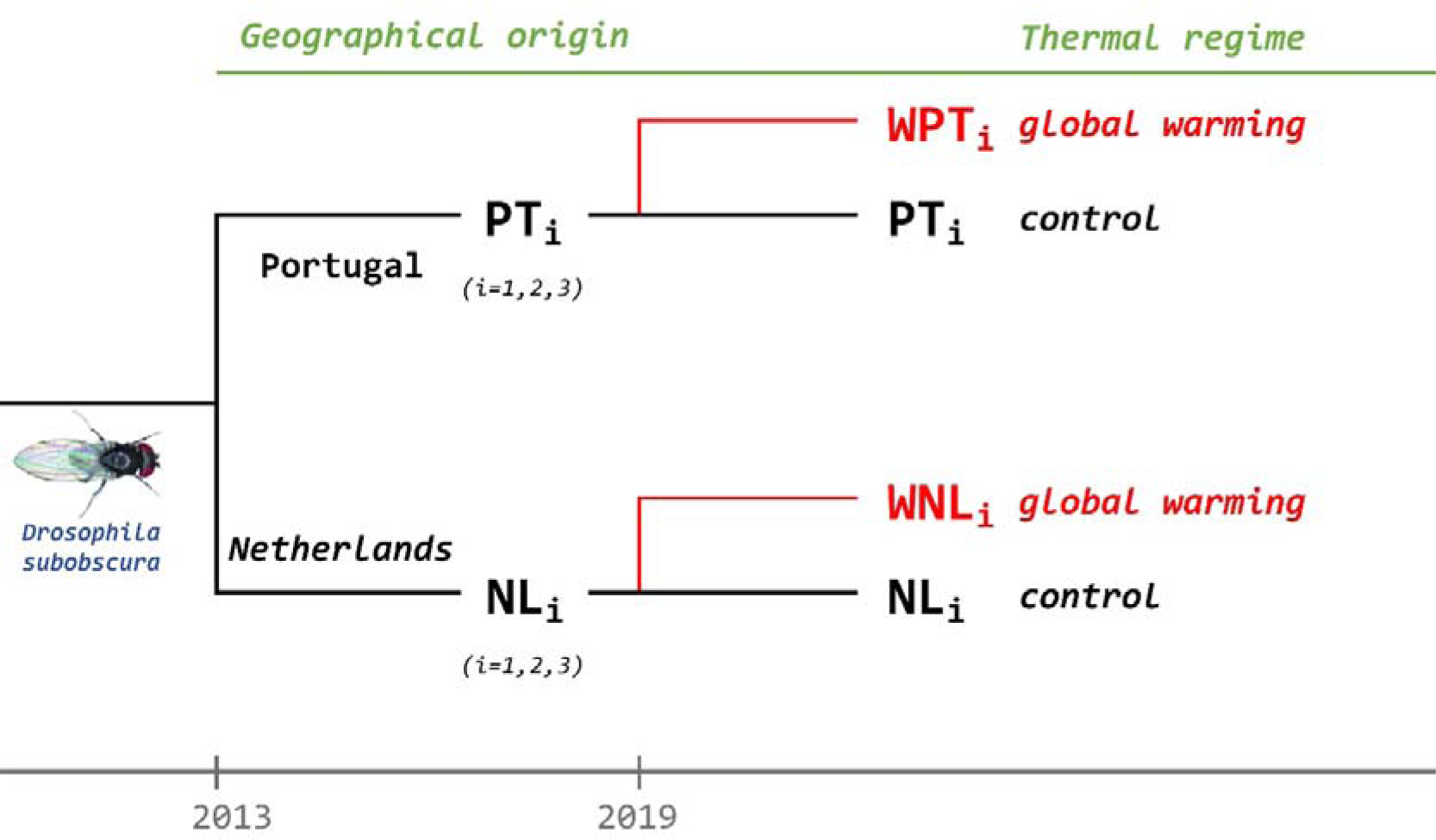
Design of the thermal adaptation experiment. Populations were collected in Adraga, Portugal (PT) and Groningen, Netherlands (NL) in 2013. By generation 70 in the lab (in January 2019), from each NL and PT replicate population (NL1-3 and PT1-3, then on designated as Ancestral population - AP) a new replicate population was derived and moved to the warming environment (see Figure 1) originating the populations WNL1-3 and PT1-3 (e.g. WPT1 and PT1 derive from the ancestral population PT1 etc) that constitute the warming thermal selection regime.

The new, warming regime had a daily fluctuation cycle that ranged between 15 C and 21 C and increased its daily mean temperature (0.18°C) and amplitude (0.54°C, with a lower minimal and higher maximal daily temperature) every generation – see Supplementary Figure 1. The increase in mean temperature in the warming regime (0.18 per generation) agrees with the expected rate of temperature increase per decade (0.19–0.63°C, (IPCC 2023)). In addition, the ratio of increase of thermal amplitude relative to the mean increase (0.54/0.18) is also aligned with the IPCC predictions (2/1). However this variation in amplitude does not intend to mimic changes in diurnal:nocturnal amplitudes as expected in nature *per se* . The rate of environmental change we imposed was high, aiming to address the potential for thermal evolution in an ectotherm species. It is comparable to the warming rates used in other experimental evolution studies (e.g. Schou et al. 2014; Rogell et al. 2014).

All experimental populations were subjected to the maintenance protocol described above, only differing in the thermal regime imposed. Specifically, egg density and adult density (70 eggs and 40 adults per vial), the one-week of adult life duration and an egg-laying period of 24h for egg collection to the next generation were maintained as in the ancestral populations. However, evolution in the warming regime led to a decoupling in generation time – from 28 days to 24 days by the time of the experiment - given the faster developmental time of populations evolving under this thermal regime. This is expected based on the association between developmental time and temperature. PT and NL population We do not expect high selective pressures for faster development as the vast majority of individuals emerged within the 4-day emergence collection period we applied and could therefore potentially contribute to the next generation.

Nevertheless, by generation 22, when the peak temperature in the warming regime reached 30.2°C, there was a significant crash in adult census sizes due to high juvenile mortality for all warming populations, which also occurred at generation 24 (average sizes of 200 and 250 respectively). As a result of this, we had to reverse the warming cycle back to the cycle of generation 20 (Santos et al. 2023b). This cycle, with a mean temperature of 21.4°C and a fluctuation between 13.5°C to 29.4°C, was then maintained till the end of the study and was the warming environment applied in phenotypic assays (see Supplementary Figure 1 and also details below). Apart from the population crashes referred above, very low population census sizes were only observed by generation 44 that motivated a two-generation transfer of the warming populations to control conditions to allow for population recovery. Otherwise, population sizes were generally high throughout the study (between 600-1000 individuals) – see Table S1.

### Experimental assays

This study includes data from the reproductive success of warming populations analysed at generations 39 and 52 of thermal adaptation. Data from the assay performed at generation 39 was already published in Santos et al. (2023b). In both assays, an orthogonal design was applied with Warming and Control populations being tested in each (Warming and Control) environment.

The assay of generation 39 involved twenty pairs of flies – placed in individual glass vials - per replicate population and environment. We used an experimental design with blocks, with each block corresponding to all same numbered replicate populations representing each Regime and History (e.g. block 1 included NL1 and WNL1, as well as PT1 and WPT1). Several racks were required in each block to cover all corresponding samples, and a pseudo-randomized approach was implemented, each rack having a subsample of each population of the corresponding block, with vials aligned per replicate population but with the relative positions changing between racks. In total, 480 pairs of flies were studied in this assay (20 pairs × 3 replicate populations × 2 thermal selection regimes (control vs warming selection regimes) × 2 historically differentiated populations (low vs high latitude) × 2 test environments (warming and control test environments) – see also (Santos et al. 2023b). At generation 52, an additional assay was performed involving sixteen pairs of females and males per replicate population and environment, in a total of 384 pairs of flies (16 pairs × 3 replicate populations × 2 selection regimes × 2 historically differentiated populations × 2 test environments).

In both assays, we estimated reproductive success as the total number of offspring derived from eggs laid by each mating pair at day 9 of adult age (a 24-hour laying period), that emerged during a 10-day period starting from the first day of emergence (maintained in the same test environment as the assayed parental generation). The ninth day of adult life was chosen as it is within the interval of the age of individuals contributing to the next generation during our population maintenance system (∼ 6 to 10 days of age). Maternal environmental effects were minimized by maintaining all assayed populations for one full generation in the control environment prior to the assay.

### Statistical methods

Data from each mating pair was used as raw data in the analyses. These included Linear mixed effects models, defining a “sum to zero” contrast option for each factor. These are orthogonal contrasts, in which the intercept refers to the average treatment effect and differences are estimated between each of the treatment means (levels) relative to the average treatment mean. Generalized linear mixed-effects models (GLMM) were applied on the whole dataset and tested with different distributions - poisson, quasipoisson, and negative binomial. To account for zeros in our dataset, zero-inflated and Hurdle models were also tested. The best overall model, based on the lowest values of Akaike information criterion (AIC), was the one assuming a quasipoisson distribution with the inclusion of a parameter accounting for zero inflation. This distribution was used in all analyses. Significance levels were obtained by applying Type III Wald chisquare tests.

We applied two overall models to the data of assays from generation 39 and 52, that varied in the random factor defined (for simplicity interactions with random factors are not presented but were included):

1. *Y = μ + History + Selection + Environment + Generation + AP{History} + History × Selection + History × Environment + History × Generation + Selection × Environment + Selection × Generation + Environment × Generation + History × Selection × Environment + History × Selection × Generation + History × Environment × Generation + Selection × Environment × Generation + History × Selection × Environment × Generation + ε*
2. *Y = μ + History + Selection + Environment + Generation + Block + History × Selection + History × Environment + History × Generation + Selection × Environment + Selection × Generation + Environment × Generation + History × Selection × Environment + History × Selection × Generation + History × Environment × Generation + Selection × Environment × Generation + History × Selection × Environment × Generation + ε*

In model (1), AP{History} is a random factor, that represents the Ancestral population (NL1-3; PT1-3) nested in the fixed factor History (PT low latitude vs NL high latitude). Thus, this term accounts for the variation between replicate populations considering the pairing due to their shared ancestry (e.g. NL1 is the ancestral population of NL1 and WNL1, nested in NL origin, etc.). Block is a random effect in model (2), being the set of same-numbered replicate populations assayed in the same pseudo-randomized experimental rack. Y is the trait under study, reproductive success. Selection is a fixed factor corresponding to the two thermal selection regimes (Warming and Control selection regimes), Environment is the fixed factor Environment representing the test environments used in the assays (Warming and Control test Environments), and Generation is the fixed factor corresponding to the two generations assayed (39 and 52). All other terms in the model represent the interactions between the fixed factors. Model (1) – ancestral population as random factor - was chosen based on lower AIC values. Considering the complexity of the model, we decided to simplify it by removing non-significant higher-order (four and three-way) interactions of fixed factors as well as their corresponding interactions including random factors. We then compared AIC values of the overall and the simplified models (see the simplified model in Table 1 and the complete model in Table S2), testing for increased fit. Considering the lower AIC values of the simplified model, we chose to interpret this model in the results section. Considering the results of the simplified model, namely the highly significant Generation term and the marginally significant Selection x Generation interaction we performed additional separate analyses for the data of each generation (39 and 52).

**Table 1.** Analysis of local adaptation across generations, populations of distinct history and environments.

| Model parameters | Estimate | St. error | Df | $\chi^2$ | p-value |
| --- | --- | --- | --- | --- | --- |
| **Intercept** | **3.185** | **0.022** | **1** | **20709.2** | **<0.001** |
| History | 0.005 | 0.022 | 1 | 0.052 | 0.821 |
| **Selection** | **-0.057** | **0.018** | **1** | **10.540** | **0.001** |
| **Environment** | **0.319** | **0.018** | **1** | **313.58** | **<0.001** |
| **Generation** | **0.087** | **0.018** | **1** | **23.907** | **<0.001** |
| History x Selection | -0.010 | 0.015 | 1 | 0.402 | 0.526 |
| History x Environment | -0.017 | 0.018 | 1 | 0.875 | 0.349 |
| **Selection x Environment** | **0.068** | **0.018** | **1** | **15.021** | **<0.001** |
| History x Generation | -0.017 | 0.016 | 1 | 1.106 | 0.293 |
| Selection x Generation | 0.026 | 0.015 | 1 | 2.917 | 0.088 |
| Environment x Generation | 0.025 | 0.018 | 1 | 1.939 | 0.164 |

Note: Total sample size: 915 observations. "Df": the degrees of freedom. Chisquare tests ( $\chi^2$ ) are presented. Statistically significant terms are represented in bold ( $p < 0.05$ )

All analyses and figures were done in R v4.3.3, with glmmTMB, car and ggplot2 packages (Wickham 2016; Brooks et al. 2017; Fox and Weisberg 2019).

## Results

To assess long-term adaptation in our warming populations we performed an overall analysis including both assays, done at generations 39 and 52 (see Table 1 and Figure 2). We observed strong evidence for differentiation in reproductive success between thermal selection regimes (significant *Selection* factor, Table 1), with a general better performance of the warming populations relative to the controls (with the exception of the patterns for the low latitude populations in the control test environment, see Figure 2 and below). We also obtained very strong evidence of a lower performance of populations when assayed in the warming test environment (significant *Environment* factor, see Table 1 and Figure 2). Importantly, we also observed that differences between thermal selection regimes changed greatly across test environments, with the warming populations showing a better performance than controls in the warming environment but less so in the control environment (significant Selection × Environment interaction; see Table 1). These differences between selection regimes across test environments were particularly evident in the case of the low latitude populations where the sign of differences reversed between environments mostly at generation 39 (see Figure 2 and below). We also found very strong evidence for variation in performance between generations, with a generally higher performance at generation 39 (significant *Generation* factor, see Table 1). Despite this, we did not find any evidence for relevant interactions of other fixed factors with generation, suggesting that the differences between generations were mostly environmental (due to asynchrony of assays) and that differences between populations of different selective regimes across environments were maintained across generations (e.g. non-significant *Selection × Environment × Generation* interaction, see Table 1). The only exception was a weak evidence for differences between selection regimes across generations (when considering both test environments) with generally higher differences between selection regimes at generation 52 than at generation 39 although this is not a conspicuous pattern (marginally significant *Selection x Generation* interaction, see Table 1 and Figure 2).

**Figure 2.**
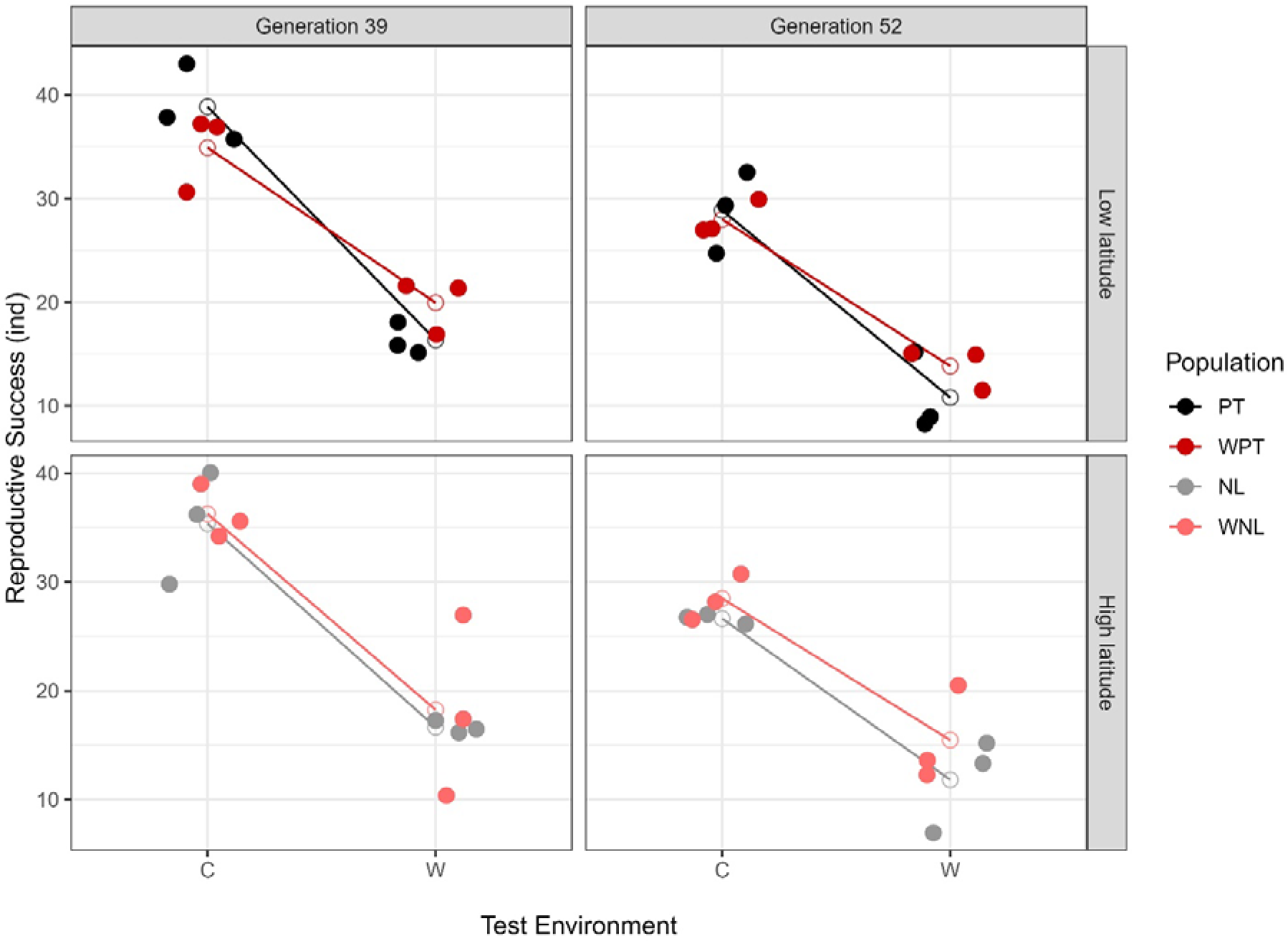
Patterns of local adaptation for low latitude and high latitude populations at generations 39 and 52. Note: Full dots represent average replicate population values, while white dots represent the averages for each combination of thermal selection regime and history (i.e. NL, WNL, PT and WPT populations). Low latitude populations are represented in the first line and high latitude in the second line; Generation 39 is represented in the first column and generation 52 in the second column. Assayed individuals were subjected to Environment C - the “Control” test environment (constant 18oC) or the Environment W - “Warming” test environment (a daily fluctuation regime with a mean temperature of 21.4°C and a fluctuation between 13.5°C to 29.4°C).

Despite the weak evidence for differences between thermal regimes across generations (see Table 1), we found advisable to analyse data from each generation separately, to address specific patterns in each generation while at the same time comparing results between the two assays. In general, we found a higher performance in the control test environment in both generations, as expected (significant *Environment* factor, χ2 = 163.10, p < 0.001 for generation 39, χ2 = 128.03, p < 0.001 for generation 52; see Table S3). Also in both assays we found strong evidence for differences in the relative performance of thermal selection regimes across test environments, with generally higher differences between warming and control populations in the warming test environment than in the control one (significant Selection x Environment interaction, χ2 = 7.540, p = 0.006 for generation 39, χ2 = 7.249, p = 0.007 for generation 52; see Table S3). In contrast, a moderate evidence for historical differences in the effect of selection across test environments was only found at generation 39 (significant *History x Selection x Environment* interaction, χ2 = 3.851, p = 0.049; see Table S3). Specifically, this was due to the pattern observed in the low latitude populations at generation 39, with the sign of differences between warming and control populations changing across test environments (Warming > Control populations in the Warming test environment; Warming < Control populations in the Control test environment). This effect, already reported in Santos et al (2023), was not observed at generation 52 (non-significant *History x Selection x Environment* interaction, see Table S3; see also Figure 2).

We estimated effect sizes for the differentiation – through Cohen’s d estimates - between thermal regimes by generations 39 and 52 (see Table S4). Let us consider first the low latitude populations. In the warming test environment at generation 39, the warming regime clearly outperformed the control, with a “very large” differentiation between the thermal selection regimes (Cohen’s d of 1.67, Warming > Control populations). By generation 52 the differentiation between selection regimes in this test environment was still present but changed to “large” (Cohen’s d of 0.99, Warming > Control populations). In the control test environment, the already described pattern of inverted sign showed a “large” effect (Cohen’s d of 1.06, Control > Warming populations) when tested at generation 39 but just a “small” differentiation by generation 52 (Cohen’s d of 0.29), suggestive of an evolutionary reduction of differences between selective regimes in the ancestral environment (see Table S4). As for the high latitude populations, there was a “small” differentiation in both test environments between selective regimes at generation 39 (Warming > Control populations), with an increase in performance of the warming populations relative to the°Controls being observed at generation 52 in both test environments: a “large” differentiation between thermal regimes in the warming environment (Cohen’s d of 0.83, Warming > Control populations), and even “very large” in the control environment (Cohen’s d of 1.22, Warming > Control populations) (see Table S4).

## Discussion

Here we provide evidence for long-term adaptation to increased temperatures in *Drosophila subobscura* populations evolving in a warming environment. In fact, by generation 52 we found that our warming populations showed a large increase in reproductive success relative to the ancestral populations (controls) when assayed in their own (warming) test environment (around 30%, see also below). Our findings point to the existence of sufficient standing genetic variation to allow for an adaptive response in our populations despite the occurrence of sporadic bottlenecks associated with thermal stress during development in particular generations and the long-term maintenance of the ancestral populations in the lab. It might also be the case that some of these episodes of high stress have opened the opportunity for interactions between drift and selection (Cohan 1986; Cohan and Hoffmann 1989) and/or reshuffling of the genetic architecture with increased additive genetic variation following bottlenecks (Bryant and Meffert 1993) that may allow for further evolutionary potential and subsequent responses.

In any case, the slow response suggests that levels of genetic variability are not high in our populations. Apart from the expected low heritability in traits closely related to fitness (Mousseau and Roff 1987; Houle 1992), as well as reduced genetic variation from lower effective population sizes in some generations. Prior laboratory adaptation (Simões et al 2017) may have also contributed to a reduction of the initial standing genetic variation. Nevertheless, we expect considerable genetic variability remains in our populations considering the genomic data from other populations frame in our lab, with similar maintenance procedures and time frame (Seabra et al. 2018).

It is important to note that the adaptive responses were only observed after the thermal increase per generation in the warming cycle was halted by generation 24, as a similar experimental test done at generation 22 failed to find evidence for thermal adaptation (Santos et al. 2023b). This finding agrees with those of another evolution experiment addressing adaptation to incremental warming in seed beetles (Hallsson and Björklund 2012; Rogell et al. 2014). These results are in accordance with the expectation that evolution lags behind environmental change, with such decoupling becoming larger with increasing rates of environmental change. Such high rates of environmental change are challenging for Evolutionary rescue (Gomulkiewicz and Holt 1995), further hindered by expected reductions in population size and in levels of genetic variation (Burger and Lynch 1995; Bell and Gonzalez 2011; Bell 2017; Martin et al. 2023). The rate of environmental change has been shown to be a critical factor for evolutionary rescue in experimental studies in bacteria, with more adaptive responses observed under slower environmental change (Liukkonen et al. 2021) and particular genotypes not being evolutionarily accessible under rapid environmental change (Lindsey et al. 2013). Consistent with this, experimental studies in ectotherms addressing short term evolution under continuously, fast increasing temperatures have shown a lack of adaptive response (Schou et al. 2014; Kinzner et al. 2019; van Heerwaarden and Sgrò 2021). We here show that adaptive responses can arise after prolonged evolution in a thermal environment with daily fluctuation and wide thermal extremes but likely not when populations face a sustained, fast rate of environmental perturbation. Consequently, this raises concerns about the ability of populations to respond to sudden thermal shifts and in particular to sustained high rates of environmental change.

We found that adaptation to the warming conditions was accompanied by changes of different magnitude in the ancestral environment, with differences between warming and controls being generally higher in the warming environment than in the ancestral environment as expected. (Santos et al. 2023b) In this context, we observed that the patterns of the two geographical populations were different, with relevant evidence that the difference in performance between the ancestral and the new, warming environment was more substantial in the low latitude warming populations than in the high latitude ones. Specifically, the low latitude warming populations showed a large decline in performance in the ancestral environment at generation 39,which was not the case for the high latitude populations. This pattern is suggestive of costs of adaptation in these populations (see Santos et al. 2023) Such adaptive costs could be due to genetic trade-offs, with alleles leading to opposite fitness effects across environments reinforcing local adaptation (Kawecki and Ebert 2004). Nevertheless, by generation 52 such differences in the ancestral environment were rather small, suggesting that such costs of adaptation were not particularly relevant. This is not a surprising finding considering other studies in the literature (see (Hereford 2009; Bono et al. 2017) for a review). It might be the case that evolution under fluctuating (heterogeneous) environments, as we use here, is less likely to allow for measurable costs of adaptation than homogeneous environments since in the latter case organisms are not exposed to different environmental conditions and consequently selection will not act to adjust/increase performance in a wider range of environments (Bono et al. 2017). Our finding of an adaptive response with transient costs suggests a possible advantage in future climatic scenarios where warmer seasons are projected to be warmer (IPCC 2023) but, at the same time, populations will need to keep their ability to cope with lower temperatures during colder seasons. However, this is most likely not enough to ensure population persistence, given the pace of the evolutionary response of the populations under study.

Even though populations of distinct history showed a quite comparable adaptive response in the warming environment at the later generation assayed (i.e. higher latitude populations showed a 31% increase relative to controls while a 28% increase was found for the lower latitude populations), we found differences between them in the tempo of evolution. In fact, the high latitude populations showed a delayed response to thermal selection, only evident after more than 50 generations of thermal evolution. In contrast, a response had been already observed in the low latitude populations by generation 39 (Santos et al. 2023b). It is an open question whether the different evolutionary dynamics and patterns observed between populations (still) reflect the geographical differentiation of this species in nature, given they had been already evolving in the control environment for around 70 generations by the time the new warming regime was imposed. Regardless, it is interesting to note that these populations still maintained clear genomic and karyotypic differentiation after the stronger phase of laboratory adaptation (∼30 generations, see Barreira 2023 and Simões et al. 2017). This suggests that some signature of past geographical history still remains. Other studies in this species have found signs of geographical differentiation associated with thermal responses (Castañeda et al. 2015; Porcelli et al. 2017). However, patterns are contrasting across traits and populations studied, with higher thermal fertility in lower latitude European *D. subobscura* populations (Porcelli et al. 2017), while the opposite trend was observed for thermal tolerance (critical thermal maximum) in South American populations (Castañeda et al. 2015). All told, more robust studies of the impact of historical backgrounds would need additional sampling of recently founded lab populations from varying latitudes.

In summary, we found that long-term adaptation to higher temperatures can occur *in D. subobscura* but the pace of such response is slow and likely dependent on low rates of environmental change. Thus, this finding supports the evidence that ectotherms may have limited capability to respond evolutionarily to temperature shifts (Kellermann and van Heerwaarden 2019) and may struggle to show an adaptive response to a fast-paced global warming. As expected, we observed more robust changes in the warming than in the ancestral environment. We further observed that the long-term evolutionary dynamics differs between populations of distinct geographical origin, with the higher latitude populations showing an adaptive response to the warming environment only in a more advanced generation (cf. (Santos et al. 2023b). This study reinforces the need for long-term evolution experiments that assess the evolution of performance under both the ancestral and the novel environment, and that scrutinize several populations to account for variation within a species. Both these components will allow for more robust assessments of species’ responses to sustained climate change.

## Data, scripts, code, and supplementary information availability

Raw data and scripts are available online at https://doi.org/10.6084/m9.figshare.25431880.v2 Simões, Pedro (2024). Complete dataset and scripts for manuscript entitled: “Long-term dynamics of adaptation to a warming environment is dependent on historical background”; Authors: Marta A. Antunes, Afonso Grandela, Margarida Matos & Pedro Simões (2024). figshare. Dataset. https://doi.org/10.6084/m9.figshare.25431880.v2

Supplementary information is available online: https://www.biorxiv.org/content/10.1101/2024.02.15.580572v5.supplementary-material

