## Supplementary Figure 1 for "Long-term evolution experiments fully reveal the potential for thermal adaptation"

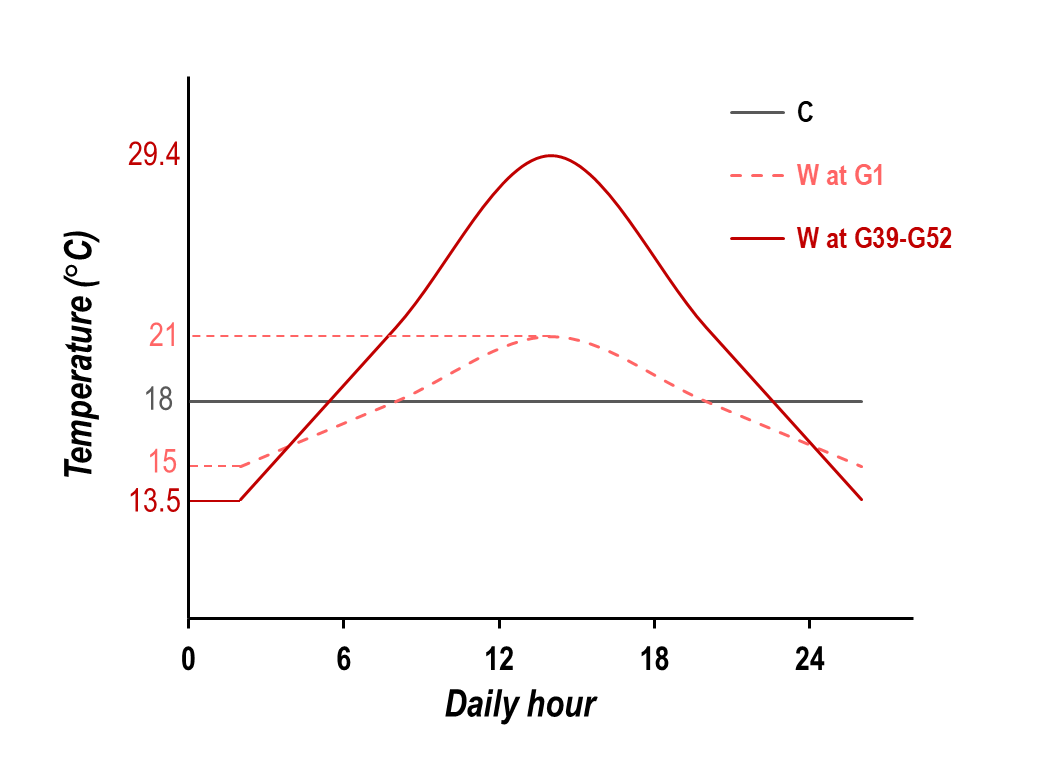

Figure S1 – Daily thermal profiles of the thermal selection regimes. “C” represents the daily thermal profile of the control regime. “W at G1” represents the thermal profile of the warming selection regime at the first generation, while “W at G39-G52” represents the thermal profile of the warming selection regime by generations 39 and 52 (maintained unchanged since generation 24 of thermal adaptation).
